# Contrasting tempos of sex chromosome degeneration in sticklebacks

**DOI:** 10.1101/2020.09.17.300236

**Authors:** Jason M. Sardell, Matthew P. Josephson, Anne C. Dalziel, Catherine L. Peichel, Mark Kirkpatrick

**Affiliations:** Department of Integrative Biology, University of Texas at Austin, Austin, TX 78712; Institute of Ecology and Evolution, University of Bern, 3012 Bern, Switzerland; Department of Biology, Saint Mary’s University, Halifax, NS, Canada B3H 3C3

**Keywords:** Sex chromosomes, stickleback, *Gasterosteus*, neo-Y chromosome, recombination, fish

## Abstract

The steps of sex chromosome evolution are often thought to follow a predictable pattern and tempo, but few studies have examined how the outcomes of this process differ between closely related species with homologous sex chromosomes. The sex chromosomes of the threespine stickleback (*Gasterosteus aculeatus*) and Japan Sea stickleback (*G. nipponicus*) have been well characterized. Little is known, however, about the sex chromosomes in their distantly related congener, the blackspotted stickleback (*G. wheatlandi*). We used pedigrees of interspecific crosses to obtain the first phased X and Y genomic sequences from blackspotted sticklebacks. Using novel statistical methods, we demonstrate that the oldest stratum of the *Gasterosteus* sex chromosomes evolved on Chromosome 19 in the ancestor of all three species. Despite this shared ancestry, the sex chromosomes of the blackspotted stickleback have experienced much more extensive recombination suppression, XY differentiation, and Y degeneration than those of the other two species. The ancestral blackspotted stickleback Y chromosome fused with Chromosome 12 less than 1.4 million years ago, which may have been favored by the very small size of the recombining region on the ancestral sex chromosome. Recombination is also suppressed between the X and Y over the bulk of Chromosome 12, although it has experienced little degeneration. These results demonstrate that sex chromosome evolution does not always follow a predictable tempo.

## Introduction

Sex chromosome evolution is thought to typically proceed via a stereotypical pathway (Charlesworth 1991; Charlesworth et al. 2005; Bachtrog 2006; Wright et al. 2016; Abbott et al. 2017; Vicoso 2019). First, an autosome becomes a sex chromosome when the autosome acquires a sex-determining gene. This can occur via a turnover event in which it gains a novel mutation or a translocation from the ancestral sex chromosome (Tanaka et al. 2007; van Doorn and Kirkpatrick 2007; van Doorn and Kirkpatrick 2010; Yano et al. 2012; Kikuchi and Hamaguchi 2013). Alternatively, sex chromosome formation can entail a fusion between an existing sex chromosome and an autosome (Charlesworth and Charlesworth 1980; Pennell et al. 2015; Matsumoto and Kitano 2016). Second, recombination is suppressed between the new X and Y (or Z and W) chromosomes, for example by an inversion (Rice 1987; Bergero and Charlesworth 2009; Charlesworth 2017). This non-recombining region is termed the sex determining region (SDR), while the segment that continues to recombine is termed the pseudoautosomal region (PAR). Various forms of selective interference then cause the SDR on the Y (or W) to degenerate via the accumulation of repeat elements, deletions, and pseudogenes (Rice 1994; Charlesworth and Charlesworth 2000; Graves 2006; Bachtrog 2008; Bachtrog 2013). As the SDR degenerates, the sex chromosomes become heteromorphic (i.e., different in size). Finally, the SDR expands via stepwise loss of recombination along the sex chromosome, e.g., due to sequential fixation of overlapping inversions (Lahn and Page 1999; Handley et al. 2004; Bergero and Charlesworth 2009; Zhou et al. 2014). This process results in “evolutionary strata” characterized by different degrees of XY differentiation and Y degeneration.

Rates of sex chromosome differentiation and degeneration can vary substantially. The Y chromosomes of *Drosophila miranda* and *Silene latifolia* degenerated greatly in 1 and 10 million years, or approximately 36 million and 7 million generations respectively (Bachtrog 2008; Papadopulos et al. 2015). In contrast, the X and Y chromosomes of the fugu (*Takifugu rubripes*) are at least 2 million years old (approximately 1 million generations), but differ only by a single nucleotide (Kamiya et al. 2012). Rates of sex chromosome degeneration also differ between primates and between birds, even in homologous strata (Hughes et al. 2005; Zhou et al. 2014). Few studies, however, have quantified the rates of sex chromosome differentiation and degeneration in congeners with young sex chromosomes. Those that have either relied on few molecular markers (e.g., Fujito et al. 2015) or failed to explicitly test whether the Y chromosomes in their study species are homologous or evolved independently (Darolti et al. 2019).

Demonstrating that the same chromosome pair determines sex in sister species is not sufficient to establish homology. The same chromosome can independently become sex-linked in multiple species. Alternatively, a turnover event in which a copy of the X chromosome evolves into new Y chromosome in one species will reset the clock of sex chromosome differentiation and Y degeneration while retaining the same sex-linked chromosome pair (Blaser et al. 2013). Understanding the origins and ages of the sex chromosomes of different species is therefore required to properly compare their tempos of sex chromosome evolution

Stickleback fishes (family Gasterosteidae) show extraordinary variation in sex determination (Fig. 1). They have experienced sex chromosome turnovers (Ross et al. 2009), transitions between XY and ZW sex determination (Chen and Reisman 1970; Ross et al. 2009; Natri et al. 2019), fusions between sex chromosomes and autosomes (Kitano et al. 2009; Ross et al. 2009; Yoshida et al. 2014; Dagilis 2019), and the origin of sex chromosomes by introgression (Dixon et al. 2018; Natri et al. 2019). As a result, nearly every species possesses different sex chromosomes.

**Figure 1:**
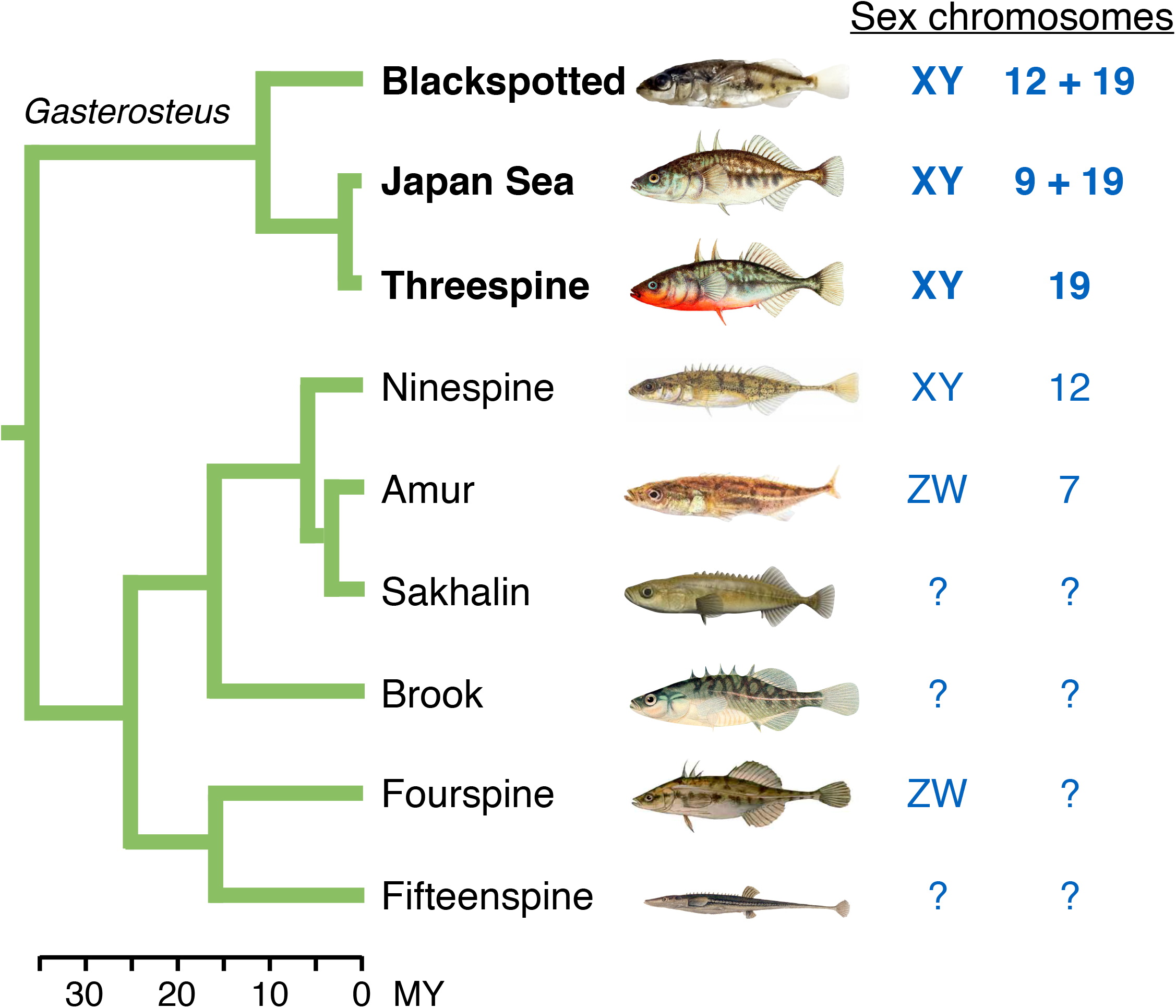
Phylogeny of several stickleback species (family Gasterosteidae) showing which chromosomes determine sex. Species in the genus *Gasterosteus* are highlighted in bold. Y-autosome fusions are indicated by “+”. Question marks denote species where the identity of the sex chromosome pair is unknown, but is inferred to be different from closely related species. (Figure after Ross et al. (2009) and Dixon et al. (2018).)

Despite frequent turnovers in other stickleback genera, Chr 19 determines sex in all three *Gasterosteus* species (Peichel et al. 2004; Kitano et al. 2009; Ross et al. 2009). The sex chromosomes in the threespine stickleback (*Gasterosteus aculeatus*) have been well characterized (Ross and Peichel 2008; Leder et al. 2010; Roesti et al. 2013; Schultheiß et al. 2015; White et al. 2015; Peichel et al. *in press*). They comprise a 2.5 Mb PAR and a 16 Mb SDR containing three strata, one of which is highly degenerated on the Y (Roesti et al. 2013; White et al. 2015; Peichel et al. *in press*). The closely-related Japan Sea stickleback (*G. nipponicus*) shares those three strata on Chr 19, indicating that recombination ceased in their common ancestor (Dagilis 2019). More recently, the ancestral Y (Chr 19) fused with Chr 9 in the Japan Sea stickleback. The resulting neo-Y chromosome now carries an additional 13.7 Mb SDR that has experienced little degeneration (Kitano et al. 2009; Natri et al. 2013; Yoshida et al. 2014; Yoshida et al. 2017; Dagilis 2019).

Much less is known about sex chromosomes in the blackspotted stickleback (*G. wheatlandi*). Using cytogenetics, Ross et al. (2009) showed that the Y chromosome comprises a fusion between Chr 19 and Chr 12 (Ross et al. 2009). They also identified nine sex-linked microsatellite markers on these two chromosomes. The Y-linked alleles of all five markers on Chr 19 did not amplify, suggesting that a wide-region of the SDR on this chromosome has degenerated. Successful amplification of markers on Chr 12 suggests that chromosome became sex-linked more recently. The blackspotted stickleback, however, has not been previously studied at the genomic level. Thus, it remains unclear whether Chr 19 was the sex chromosome in the common ancestor of all *Gasterosteus*, or whether it independently evolved into a sex chromosome in multiple species. Chr 12 also determines sex in the more distantly-related ninespine stickleback (*Pungitius pungitius*), and is another candidate for the ancestral sex chromosome in *Gasterosteus* (Ross et al. 2009; Shapiro et al. 2009; Dixon et al. 2018; Natri et al. 2019).

In this paper, we present the first genomic investigation of the blackspotted stickleback sex chromosomes. Using phased genome sequences obtained from pedigreed crosses, we find that the blackspotted stickleback X and Y are highly differentiated along nearly the entire length of Chr 19, and that the entire SDR on Chr 19 exhibits extreme Y degeneration. This situation is in stark contrast to that in the threespine and Japan Sea sticklebacks, where extreme degeneration is limited to the oldest stratum on Chr 19. The fused neo-Y of Chr 12 in blackspotted stickleback also contains a large SDR that shows relatively little degeneration. This fusion occurred recently and independently of its recruitment as a sex chromosome in ninespine stickleback. We conclusively demonstrate homology between the ancestral blackspotted and Japan Sea stickleback Y chromosomes (Chr 19) using two approaches: a novel method for detecting shared duplications onto the Y, and a gene-tree based approach. Thus, the extensive differentiation between the blackspotted X and Y evolved over the same time scale as the more limited differentiation seen in the threespine and Japan Sea sex chromosomes. We conclude that evolution of young sex chromosomes does not always follow a predictable mode or tempo even when species share the same ancestral sex chromosome.

## Results

We obtained phased sequences of X and Y chromosomes from 15 interspecific crosses, each consisting of a blackspotted stickleback father, a threespine stickleback mother, one daughter, and one son. All four members of each family were shotgun sequenced, and the haploid genome sequences of the blackspotted father’s X-bearing and Y-bearing sperm were determined from patterns of transmission (see also Sardell et al. 2018; Dagilis 2019). In this way, we sequenced 15 independent blackspotted X chromosomes and 15 independent blackspotted Y chromosomes. All reads were mapped to the repeat-masked version of the threespine stickleback reference genome (Glazer et al. 2015). This reference was produced from a female and so lacks a sequence for the Y. We did not map reads to the recently-published threespine stickleback Y reference sequences (Peichel et al. *in press*) for reasons described in the ‘<u>Sequence assembly & SNP calling</u>’ section of the Materials and Methods. All genome positions given for data from the blackspotted stickleback refer to the threespine reference.

SDRs often show two patterns (Vicoso and Bachtrog 2015; Palmer et al. 2019). First, lack of recombination allows the X and Y to accumulate lineage-specific mutations, leading to differentiation between them, as seen (for example) by elevated *F*_ST_. Second, as the sequences of the X and Y continue to diverge, some Y-linked reads will not map to the X chromosome reference, particularly when deletions or repeat elements accumulate on the Y. Consequently, the mean read depth across the sex chromosomes will be lower in males than in females, and the ratio of read depths in males vs. females (hereafter termed “read depth ratio”) will decrease.

SDRs also exhibit a property that we call XY monophyly (Dixon et al. 2018; Toups et al. 2019). If fixation of an inversion on the Y causes the SDR to expand, lack of recombination between the X and Y causes all Y sequences and no X sequences within the SDR to descend from a single ancestor’s chromosome in which inversion first occurred. That is, gene trees will show the Ys falling within a monophyletic clade with respect to the Xs. The Xs will also be reciprocally monophyletic if enough time has passed for them to coalesce since the inversion fixed. The argument works conversely when an inversion is fixed on the X, and it applies equally to any mechanism that completely blocks recombination between the X and Y. In gene trees from the PAR, historical recombination in the sampled population causes sequences from the Xs and Ys to be intermingled.

### The SDR on the ancestral sex chromosome (Chr 19)

Our results for Chr 19 show that the SDR spans nearly the entire chromosome, that the X and Y are highly differentiated, and that the Y is highly degenerate (Figs. 2 and 3). More SNPs have a read depth ratio nearer to 0.5 than to 1.0 on Chr 19 (Fig. 2A). This result is commonly interpreted as a signal of extensive Y degeneration (e.g., Roesti et al. 2013; Vicoso and Bachtrog 2013; Zhou et al. 2014; Darolti et al. 2019; Palmer et al. 2019). It indicates that many regions present on the X have been deleted from the Y and/or that many reads from the Y have diverged from the X to the point where they fail to map to the X reference scaffold. Plotting the read depth ratio along the chromosome demonstrates that the highly degenerate region of the SDR spans nearly all of Chr 19 (Fig. 2B). Mean *F*_ST_ between the X and Y is also highly elevated over nearly the entire chromosome (Fig. 3A). XY monophyly confirms that much of Chr 19 is non-recombining (Fig. 3B). Among the 100 Kb windows in the SDR, 74% (150/203) exhibit X monophyly, as expected if recombination between the X and Y ceased long ago or if an inversion recently swept to fixation on the X. Patterns of Y monophyly are noisier, with 46% (94/203) of windows in the SDR exhibiting monophyly. Windows exhibiting incomplete Y or X monophyly within the SDR likely reflect genotyping and phasing errors in regions with high Y degeneration, as hemizygosity in males causes SNP calling tools and phasing algorithms to erroneously impute X alleles onto the Y (see Materials and Methods). We applied several bioinformatic filters to discard hemizygous regions, but some may still be represented in our data set. Windows with incomplete Y monophyly may also have resulted from rare gene conversion events.

**Figure 2:**
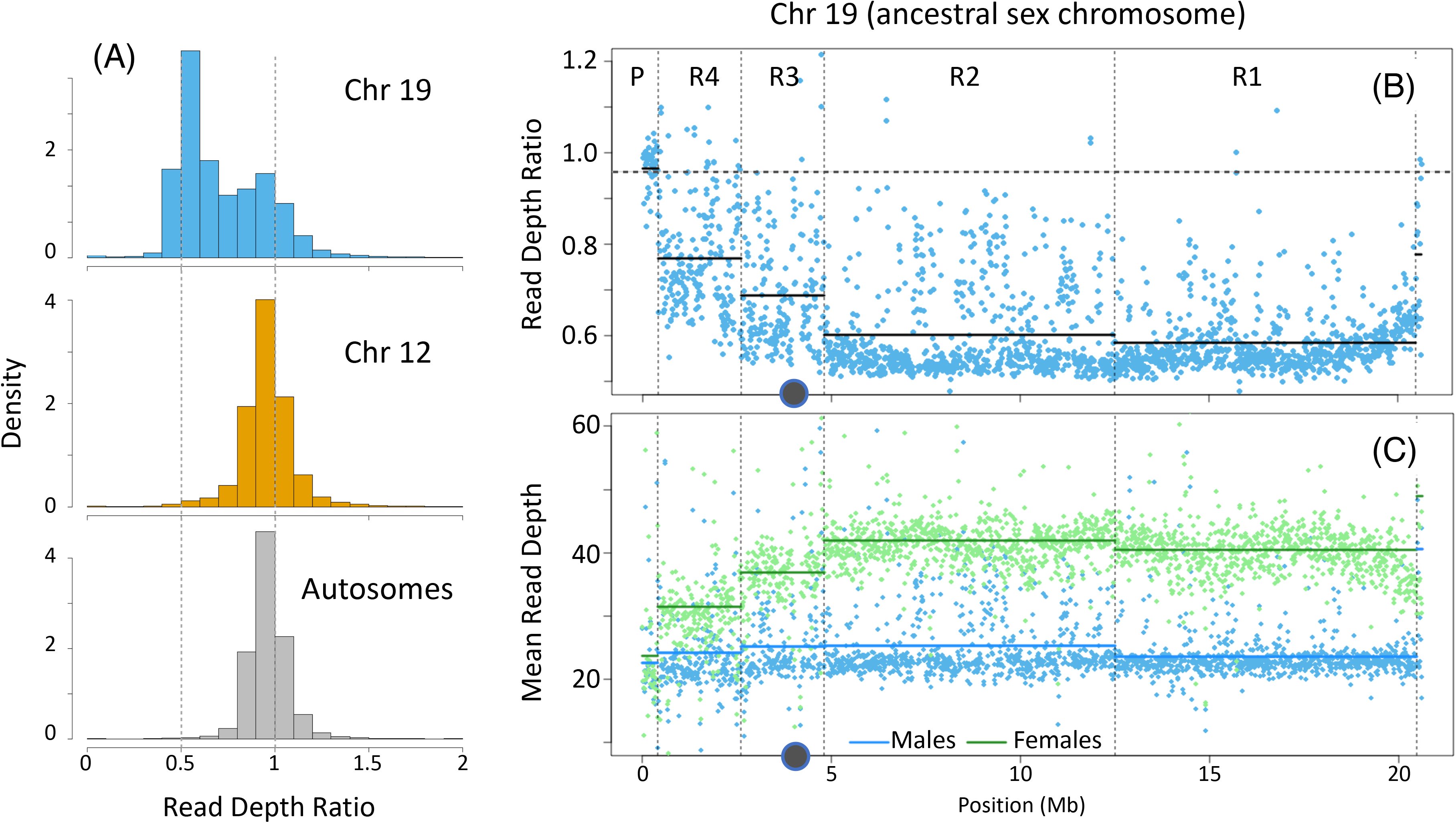
Read depth statistics for 15 sons and 15 daughters from the pedigrees. (A) Histogram of male/female read depth ratio for all SNPs on the two pairs of sex chromosomes and the autosomes. Dashed vertical lines show values expected with Y chromosomes that are highly degenerated (ratio = 0.5) and non-degenerated (ratio = 1). (B) Plot of mean male/female read depth ratio along Chromosome 19. Dashed horizontal gray line represents autosomal mean. (C) Plot of mean read depth in sons (blue) and daughters (green) along Chromosome 19. Dots are averages in 10 Kb windows. Dashed vertical lines represent boundaries between the PAR (labeled P) and the four regions in the SDR (labeled R1 to R4) that were identified using methods described in text. Solid horizontal lines show the means for the regions. Gray dots underneath the plots denote the location of the centromere on the X chromosome in threespine stickleback.

**Figure 3:**
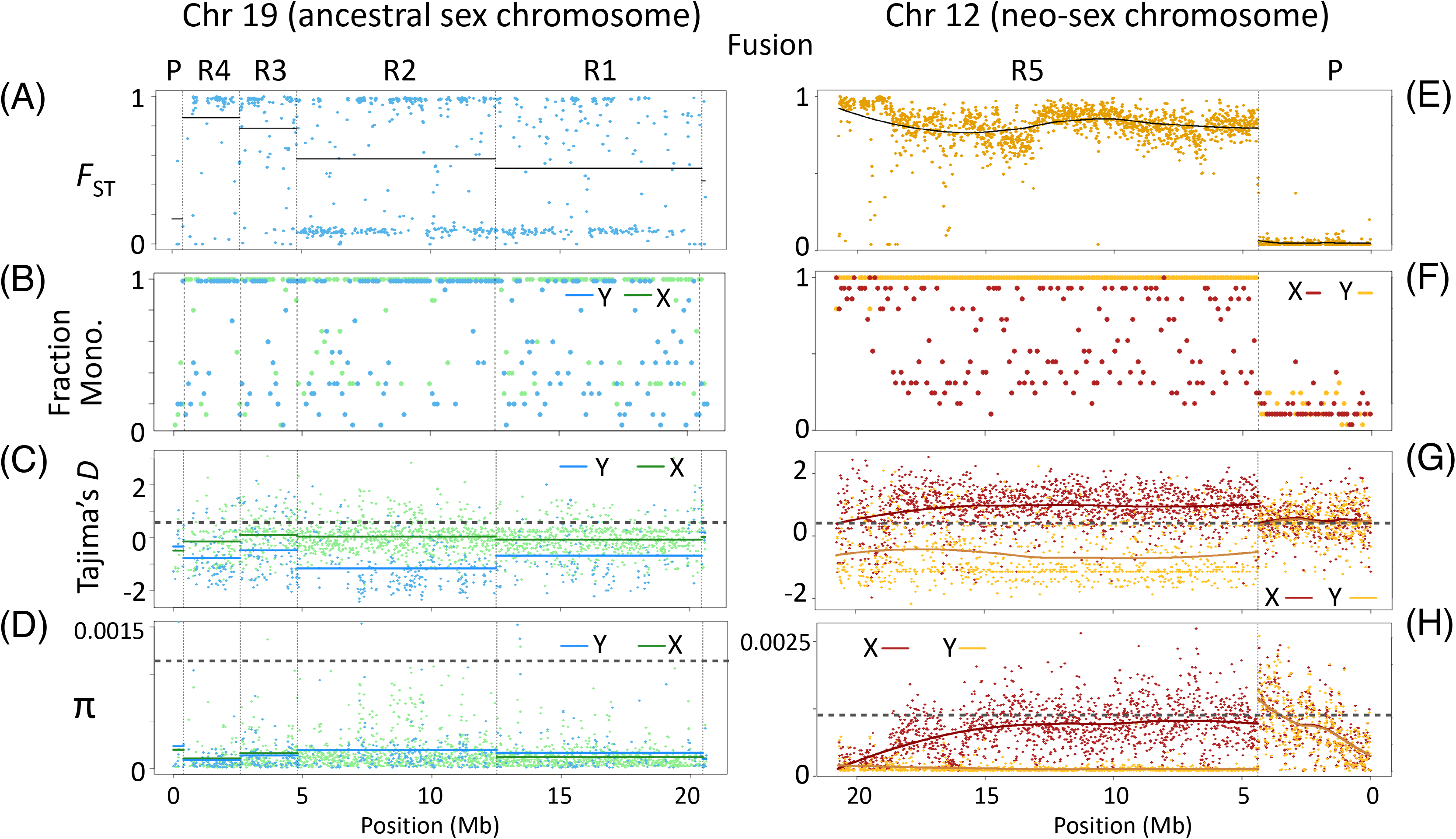
Population genetic statistics for 15 X and 15 Y chromosomes in blackspotted stickleback. Points are averages in 10 Kb windows. Panels A-D represent the ancestral sex chromosome (Chr 19) and Plots E-H represent the fused neo-sex chromosome (Chr 12). (A,E) *F*_ST_ between X and Y chromosomes. (B, F) Fraction of Y (or X) chromosomes that fall within the largest monophyletic clade of Y (or X) chromosomes on the gene tree. (C, G) Tajima’s *D* across X and Y chromosomes. (D, H) Genetic diversity (π) across X and Y chromosomes. Solid lines show the means for the PAR (P) and the four regions (R1 to R4) on the SDR of Chr 19. Loess best-fit curves show the means for the PAR (P) and SDR (R5) of Chromosome 12. Horizontal gray lines in panels C, D, G, and H show autosomal means.

A small region (400 Kb, or about 2% of the chromosome) at the end of Chr 19 distal to the fusion is a PAR. Read depth between sons and daughters is nearly equal here (Fig. 2B), and most windows have low *F*_ST_ between the X and Y (Fig. 3A). The gene trees in this region do not exhibit X or Y monophyly, indicating that it continues to recombine (Fig. 3B).

Read depth ratios vary along the SDR on Chr 19, which may indicate the presence of strata (Fig. 2B). We identified putative strata using an algorithm that detects changepoints in the read depth ratio data (Killick and Eckley 2014). This method groups Region 1, which spans from 12.5 Mb to the fusion and corresponds to the oldest stratum in threespine stickleback (Roesti et al. 2013; White et al. 2015; Peichel et al. *in press*), with Region 2 (4.8 - 12.5 Mb). Both show extensive Y degeneration, having mean read depth ratios close to 0.5, and are only revealed to be different strata by multi-species gene trees (see below). The read depth ratios in Regions 3 (2.6 - 4.8 Mb) and 4 (0.4 - 2.6 Mb) are reduced to a lesser degree. Another pattern emerges, however, when we examine males and females separately: the read depth is notably less in Regions 3 and 4 relative to Regions 1 and 2 in females, but consistently low in males (Fig. 2C). This is surprising because Y degeneration should only affect read depths in males.

Genomic divergence between the X and Y at synonymous site (*d*_S_) also varies between putative strata on Chr 19, as expected if they stopped recombining at different times (Supp. Fig. S1). Notably, genes in Region 4 have significantly lower mean *d*_S_ than genes in Regions 1, 2, and 3 based on Mann-Whitney *U* tests (*p* ≤ 10^−13^). Genes in Region 3 have significantly lower *d*_S_ than genes in Regions 2 (*p* = 10^−6^), but not lower than Region 1 after accounting for multiple comparisons (*p* = 0.01). Genes in Region 2 have significantly higher *d*_S_ than genes in Region 1 (*p* = 0.0005). Genomic divergence between the X and Y at nonsynonymous sites (*d*_N_) exhibits similar patterns across these regions (Supp. Fig. S2A), with statistically significant differences between all regions (*p* < 10^−6^) except Regions 1 and 2 (*p* = 0.03). The *d*_N_*/d*_S_ ratio between the Xs and Ys is similar across most regions (Supp. Fig. S2B), and only genes in Regions 1 and 3 are significantly different after controlling for multiple comparisons (0.90 vs. 1.13, *p* = 0.001).

We estimated the ages of the putative strata by comparing *d*_S_ between the blackspotted stickleback X and Y to *d*_S_ between the blackspotted and threespine stickleback X chromosomes in each region, using 14.3 million years as the estimated date of the speciation event (Varadharajan et al. 2019). Based on this approach, Regions 1 and 3 both stopped recombining around the time of the species split. We estimate that Regions 2 and 4 have been non-recombining for approximately 12.3 and 10.5 million years, respectively.

We calculated several additional population genetics statistics for the sex chromosomes. Molecular diversity (π) on Chr 19 is very low on both the X and Y chromosomes (Fig. 3D). Tajima’s *D* is strongly negative across the Y (Fig. 3C), suggesting that the Y recently experienced a selective sweep, consistent with the spread of the fusion that formed the neo-Y. Tajima’s *D* is close to 0 on the X (Fig. 3C), and is significantly lower than values for the autosomes (range 0.53 - 0.59). Together these results suggest that the X has also recently experienced one or more selective sweeps, while the strongly positive value of Tajima’s *D* on the autosomes implies that the species’ population size has recently decreased.

### The SDR on the neo-sex chromosome (Chr 12)

Different patterns emerge on the neo-sex chromosome, Chr 12, where an autosome has fused to the Y chromosome (Chr 19). This fusion resulted in a doubling of the size of the Y, as Chrs 12 and 19 are both approximately 20 Mb. Y monophyly clearly shows that the SDR has expanded across most of Chr 12, extending 16.4 Mb from the fusion (Region 5). Lack of complete X monophyly in most regions of the SDR indicates that recombination between the X and Y in Region 5 was recently suppressed (*i.e*., on the order of 2*N*_e_ generations ago) (Fig. 3F). The mean read depth ratio in the SDR is nearly equal to its value on autosomes, indicating that the neo-Y is young and has not degenerated much (Supp. Fig. 3). Values of *d*_S_ between the X and Y within the SDR of Chr 12 are low and indicate that a single stratum evolved less than 1.4 million years ago (Supp. Fig. S1). This is likely an overestimate for the age of the SDR because the X clade is polyphyletic with respect to the Ys across much of the SDR (Fig. 3F). Thus, the most recent common ancestor of all Y and X chromosomes predates the formation of the SDR on Chr 12. The PAR makes up the remaining 4.4 Mb of the chromosome distal to the fusion, as shown by low values of *F*_ST_ between the X and Y (Fig. 3E) and the lack of monophyly of either X or Y chromosomes (Fig. 3F). Thus, the blackspotted Y has two PARs, a small one on the end of Chr 19 and larger one on the end of Chr 12.

Population genetics statistics for the Chr 12 neo-sex chromosomes reveal recent evolution of the neo-Y and neo-X. The neo-Y SDR exhibits nearly no polymorphism (Fig. 3H) and Tajima’s *D* is strongly negative across its length (Fig. 3G). These patterns are consistent with a very recent sweep, potentially associated with the Y fusion or expansion of the SDR. Molecular diversity within the SDR of Chr 12 is much higher on the neo-X than on the X of Chr 19 and is slightly lower than the autosomes (Fig. 3H). Tajima’s *D* is larger on the neo-X than the autosomes (Fig 3G). That result is consistent with the 25% reduction in population size that it experienced after the fusion changed it from an autosome to an X.

### The Y chromosome originated in the *Gasterosteus* ancestor

The degree of X-Y divergence and Y degeneration on Chr 19 are dramatically greater in the blackspotted sticklebacks than in the other two species of *Gasterosteus* that have been studied on a genomic level (Ross and Peichel 2008; Leder et al. 2010; Natri et al. 2013; Roesti et al. 2013; Yoshida et al. 2014; Schultheiß et al. 2015; White et al. 2015; Yoshida et al. 2017; Dagilis 2019; Peichel et al. *in press*). One hypothesis is that their Y chromosomes are not homologous, and the blackspotted Y is much older. We used two approaches to falsify that hypothesis. The first is based on a new method that identifies shared genomic rearrangements, and the second uses an analysis of gene trees. The results show clearly that the oldest stratum of the Y chromosome (Region 1 on Chr 19) evolved in the common ancestor of all three *Gasterosteus* species.

#### Shared duplications onto the Y chromosome

Homology between Y chromosomes can be inferred from shared Y-specific chromosomal rearrangements. Bissegger *et al*. (2019) discovered that at least 38 small autosomal regions have been duplicated onto the Y chromosome SDR in threespine stickleback. These regions appear to have extreme differentiation (e.g., *F*_ST_) between males and females on autosomes, which is an artifact that results when sequencing reads from duplicated regions on the Y chromosome mismap to their autosomal paralogs.

Exploiting that discovery, we asked if autosome-to-Y duplicates are shared between blackspotted sticklebacks and other *Gasterosteus* species. We first calculated *F*_ST_ between paternally inherited sequences for sons vs. daughters in 10 Kb windows across all autosomes in our blackspotted stickleback pedigrees. We then calculated *F*_ST_ across the same windows for a comparable set of pedigrees involving Japan Sea stickleback males studied by Dagilis (2019). Considering those windows whose *F*_ST_ values fall in the top 2% of the distribution, we find that 183 outlier windows are shared between blackspotted and Japan Sea sticklebacks. This number is far greater than expected by chance (*n* = 14, *p* < 0.00001, chi-squared test). Of these windows, 98 have SNPs shared by both species. Three of these windows contain multiple SNPs where both species have high *F*_ST_ (> 0.25) between the X and Y and the same male-specific allele (Fig. 4). These SNPs, which we refer to as “homologous Y duplicates”, provide very strong evidence that these three regions were duplicated from the autosomes to the Y in the ancestor of blackspotted and Japan Sea sticklebacks (Fig. 4). Intriguingly, one of the windows (17.15 to 17.16 Mb on Chr 8) contains the ortholog to the putative male-determining gene (*Amhy*) in threespine stickleback, which arose by a duplication from Chr 8 to Chr 19 (Peichel et al. *in press*). This finding suggests that all *Gasterosteus* species share the same master sex determining gene.

**Figure 4:**
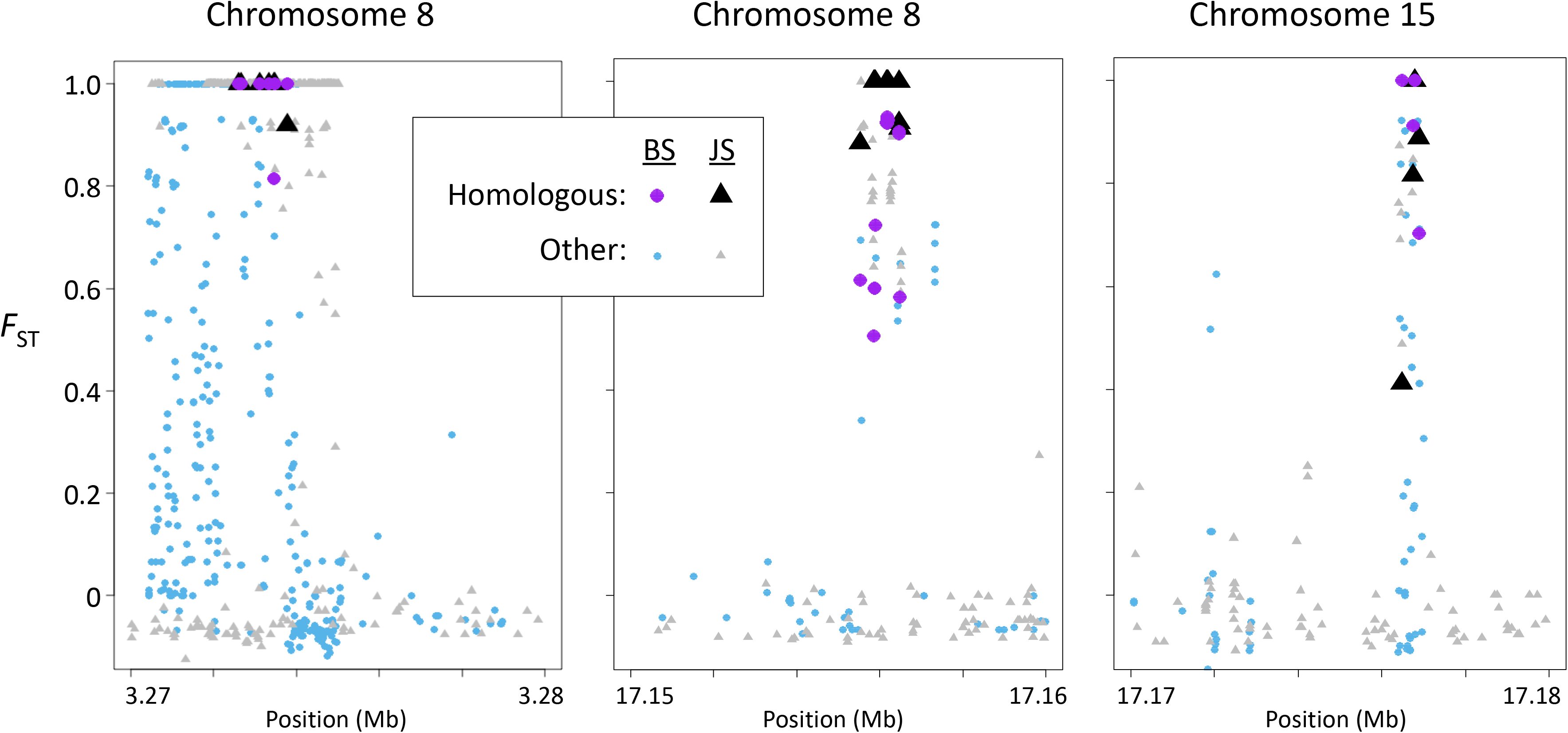
Three 10 Kb windows on autosomes include regions that duplicated onto the Y chromosome (Chr 19) in the shared ancestor of *Gasterosteus* sticklebacks. *F*_ST_ between sons and daughters for alleles inherited from the father is shown for each SNP. Circles represent SNPs in blackspotted (BS) sticklebacks and triangles represent SNPs in Japan Sea (JS) sticklebacks. “Homologous” denotes SNPs that have *F*_ST_ > 0.25 and a male-specific allele in both species, which provides strong evidence for Y chromosome homology. The window between 17.15 to 17.16 Mb on Chr 8 contains the ortholog to the putative male-determining gene (*Amhy*) in threespine stickleback (Peichel et al. 2020).

The homologous Y duplicates within these three windows show additional features consistent with autosome-to-Y duplications. Read depth is consistently higher in males than females (Supp. Fig. S4A), as expected if males have one or more Y duplicates in addition to the autosomal paralog. Reads containing the male-specific alleles for these SNPs typically comprise much less than half of the total reads mapping to the region (Supp. Fig. S4B). This pattern is expected when a mutation fixes in the Y paralog, since there are twice as many copies of the autosomal paralog in the genome. Finally, we find that the high *F*_ST_ regions in each of these three windows BLAST with high similarity to at least one region in the threespine stickleback Y reference (Peichel et al. *in press*), but not to any other region on the autosomes or X chromosome in the refence genome. All these data are strong evidence that these SNPs fall within regions that duplicated onto the Y chromosome from autosomes in the common ancestor of *Gasterosteus* sticklebacks.

#### Gene trees

Different hypotheses for the evolution of sex chromosomes predict different gene tree topologies (Dixon et al. 2018). If a non-recombining sex chromosome evolved in the common ancestor of two species, then their Y chromosomes will be more closely related to each other than to the X chromosomes of their own species (Fig. 5A). Conversely, if their sex chromosomes arose from autosomes independently, then each Y chromosome will be most closely related to the X chromosome from the same species (Fig. 5B). Finally, if the Y in one species arose from an X chromosome, then the new Y will form a clade with the X chromosome in that species, which will in turn be sister to the X chromosome from the other species (Fig. 5C).

**Figure 5:**
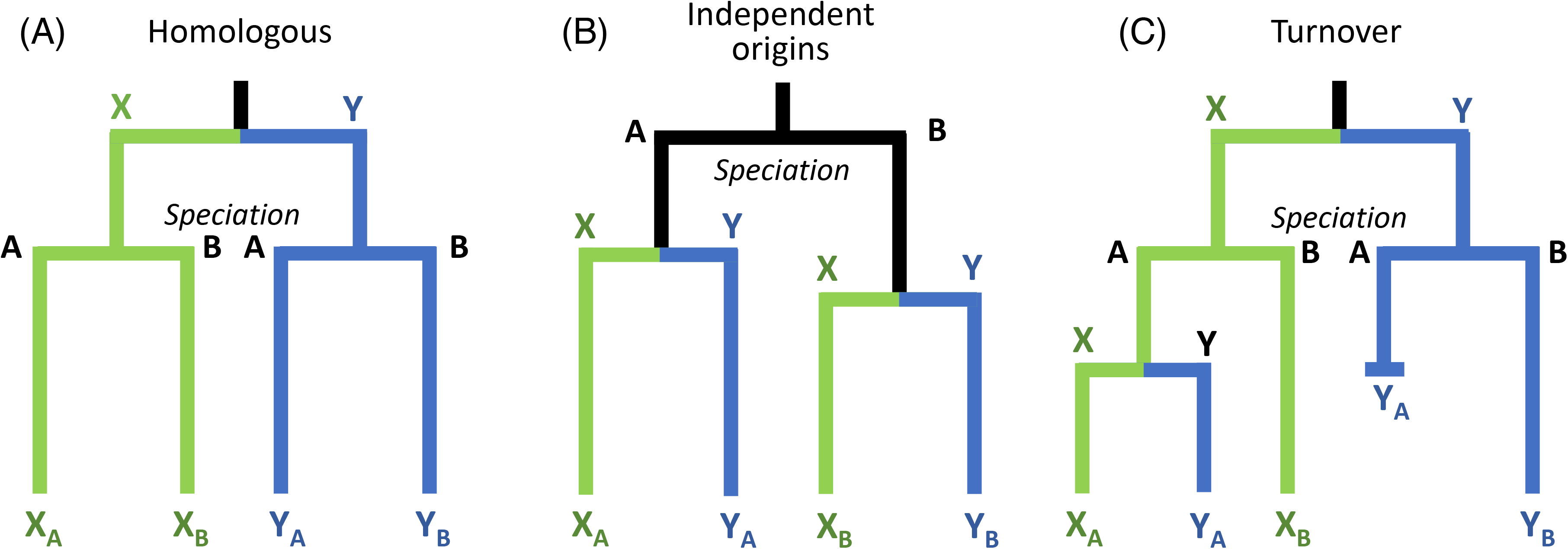
Different evolutionary histories of sex chromosomes result in different gene tree topologies. (A) Topology when the Y chromosomes of two species (A and B) are homologous, having originated in their common ancestor. (B) Topology when the Y chromosomes originated independently in two species. (C) Topology when the X and Y are retained from the ancestor in Species B, while a new Y was derived from an X chromosome and replaced the ancestral Y in a turnover event in Species A.

We determined the evolutionary history of the *Gasterosteus* sex chromosomes by constructing gene trees for non-overlapping 100 Kb windows. The sequence data come from two sets of pedigrees. From this study, we used four phased X and four phased Y chromosomes from the blackspotted stickleback fathers and eight phased X chromosomes from the threespine stickleback mothers. From a previous study by Dagilis (2019), we used four phased X and four phased Y chromosomes from Japan Sea stickleback fathers and eight X chromosomes from threespine stickleback mothers. As an outgroup, we included a Chr 19 from a ninespine stickleback that was computationally phased by Dixon et al. (2018).

Two topologies dominate the gene trees in the SDR of Chr 19 (Fig. 6). In Region 1, the most common topology is one in which the blackspotted Ys are sister to the Japan Sea Ys, and the blackspotted Xs are sister to the Japan Sea Xs. This topology, which is not found in other regions, suggests that this stratum evolved in the common ancestor of these two species and that neither species has experienced sex chromosome turnover since (compare to Fig. 5A). In Regions 2 and 3, most windows exhibit a topology in which the blackspotted Xs and Ys are sister to one another, as are the Japan Sea Xs and Ys. This topology suggests the SDR expanded into these regions independently in the two species after they diverged (compare to Fig. 5B). Region 4 lies in the blackspotted SDR, while it is in the Japan Sea PAR. As a result, the blackspotted Xs and Ys form clades that are sister to one another, while the Japan Sea Xs and Ys are intermingled (since they continue to recombine). Finally, the region from 0 to 400 Kb features clades separating the species, but no differences between the Xs and Ys within species, consistent with it being a PAR in both species. Few windows on Chr19 show topologies consistent with an X-to-Y chromosome turnover, and most of those that do fall within regions that also feature several windows with biologically implausible topologies. Thus, these patterns likely reflect genotyping and phasing errors resulting from degeneration of the Y chromosomes in one or both species.

**Figure 6:**
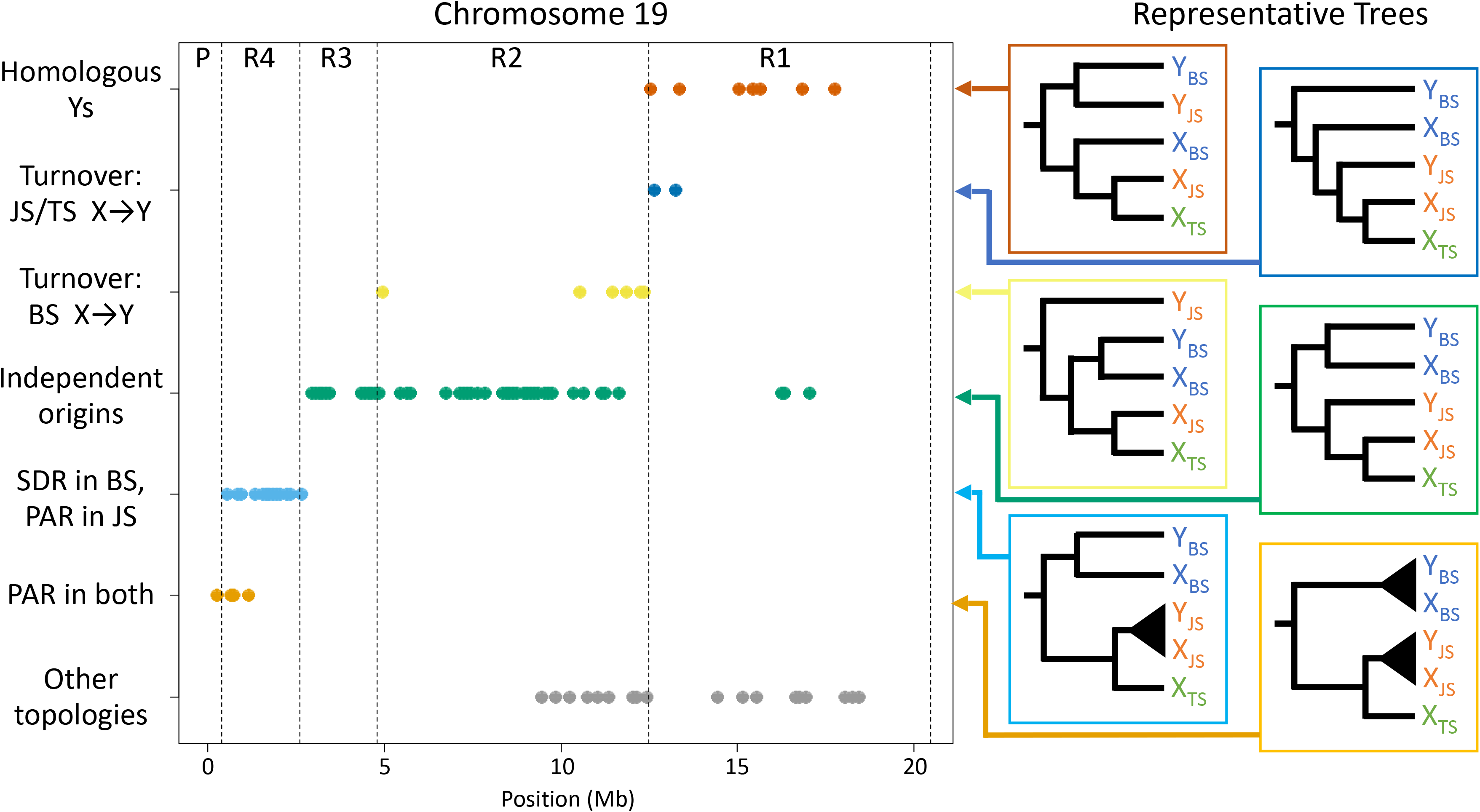
Gene tree topologies along Chr 19 reveal different evolutionary histories. Each dot represents the maximum likelihood topology for a 100 Kb window. Representative trees associated with each topology are shown at right (BS = blackspotted, JS = Japan Sea, TS = threespine). “Other topologies” indicates topologies that do not correspond to a plausible evolutionary history, and likely result from genotyping and/or phasing error. Most windows in Region 1 have a topology indicating that the stratum arose in the shared *Gasterosteus* ancestor. Most windows in Regions 2 and 3 have a topology consistent with strata that formed independently in blackspotted and in the ancestor of Japan Sea and threespine sticklebacks. Most windows in Region 4 have a topology consistent with an SDR in blackspotted and a PAR in the ancestor of Japan Sea and threespine sticklebacks.

Chr 12 is sex linked in blackspotted sticklebacks as well as the distantly related ninespine stickleback, but not in the congeneric Japan Sea or threespine sticklebacks. Gene trees across the SDR of Chr 12 in the blackspotted stickleback confirm that its neo-X and neo-Y are closely related to one another, and that its neo-Y is young since its sequences are embedded with the neo-X sequences (Supp. Fig. 5). Thus, Chr 12 has independently evolved to be a sex chromosome in the blackspotted and ninespine stickleback.

## Discussion

Many studies have analyzed variation in the mode and tempo of sex chromosome evolution, but most have compared species that share ancient sex chromosomes (e.g., Hughes et al. 2005; Goto et al. 2009; Zhou et al. 2014; Xu et al. 2019a; Xu et al. 2019b) or that have non-homologous sex chromosomes (e.g., Vicoso et al. 2013; Hough et al. 2014; Papadopulos et al. 2015; Crowson et al. 2017; Jeffries et al. 2018). Clades in which several species possess sex chromosome pairs that descend from a single recent ancestor offer unique insight into the predictability of this process. *Gasterosteus* sticklebacks present an excellent opportunity to quantify variation in the tempo of sex chromosome evolution, as we have established that the Y chromosomes of all three species are homologous. Using phased X and Y chromosomes, we obtained the first genomic characterization of the blackspotted stickleback sex chromosomes. This allows us to compare and contrast their structure and evolutionary history to that of the same pair of sex chromosomes in threespine and Japan Sea sticklebacks (hereafter called the “threespine clade”) which have been well-studied (Ross and Peichel 2008; Leder et al. 2010; Natri et al. 2013; Roesti et al. 2013; Yoshida et al. 2014; Schultheiß et al. 2015; White et al. 2015; Yoshida et al. 2017; Dagilis 2019; Peichel et al. *in press*). We found that the blackspotted stickleback has experienced more extensive suppression of recombination, sex chromosome differentiation, and Y degeneration than the threespine or Japan Sea sticklebacks experienced over the same time scales despite a common origin.

### Contrasting patterns in other sticklebacks

We observed striking differences between the sex chromosomes of the blackspotted and the threespine clade. Stratum 1 in the threespine clade, which is the oldest in that group (Roesti et al. 2013; Dagilis 2019; Peichel et al. *in press*), is also present in the blackspotted sex chromosomes (Region 1) and evolved in their shared ancestor. Patterns of divergence at synonymous sites suggests that this shared stratum formed around the speciation event between blackspotted stickleback and the threespine clade (14 million years ago). In contrast, Peichel et al. (*in press*) estimated that the oldest stratum of the threespine stickleback sex chromosomes formed closer to 21.9 million years ago. The latter estimate is probably more accurate, as it was based on a fully sequenced Y chromosome assembly. In contrast, our method likely underestimates *d*_S_ in the highly degenerate regions of the blackspotted Y due to genotyping errors. When a locus is missing from the Y, genotyping algorithms erroneously impute maternal threespine X alleles onto the phased blackspotted Y sequences. These errors bias our divergence time estimate downwards towards the time when the threespine and blackspotted X chromosomes diverged. We applied filters to remove most hemizygous errors from our sequences, but filtering also decreases estimates of pairwise diversity. Taken together, we conclude that the Y has extensively degenerated across this stratum in all species and that it has not recombined for approximately 21.9 million years, which is before the ancestors of the blackspotted and threespine sticklebacks diverged 14 million years ago (Fig. 7) (Dagilis 2019; Varadharajan et al. 2019; Peichel et al. *in press*).

**Figure 7:**
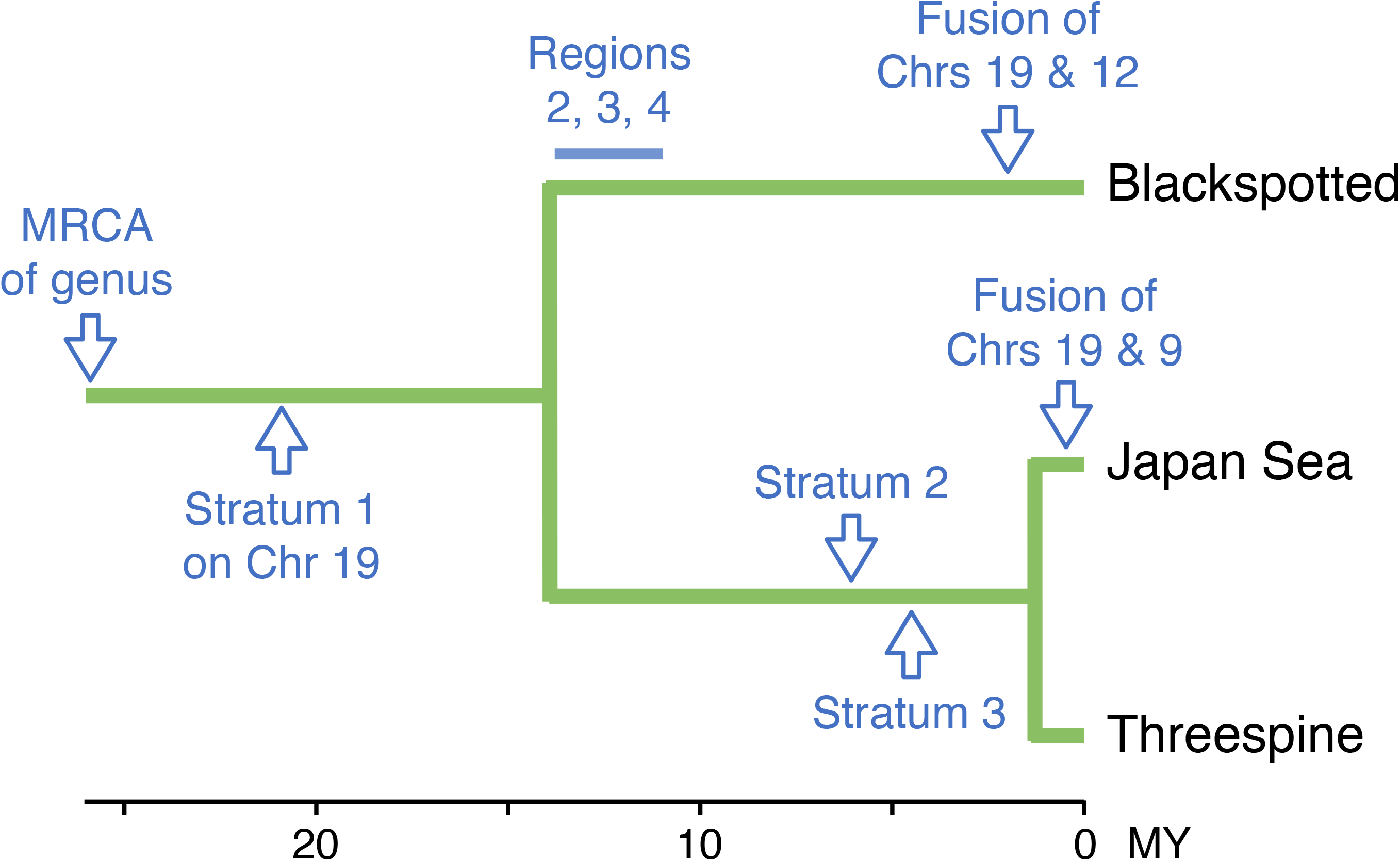
Estimated dates of the major steps in the evolution of sex chromosomes across *Gasterosteus*. The most recent common ancestor (MRCA) of *Gasterosteus* and all other stickleback species is on the left. The oldest stratum formed in the ancestor of the genus, and the SDR independently expanded across the sex chromosomes in blackspotted stickleback (Regions 2-4) and the threespine clade (Strata 2 and 3). The Y chromosome recently fused with different autosomes in blackspotted and Japan Sea stickleback. Estimated dates are from Varadharajan et al. (2019), Peichel et al. (*in press*), and Dagilis (2019).

Following the divergence of the blackspotted and threespine clade, the SDR independently expanded across most of the remainder of Chr 19 (Fig. 7). In the threespine clade, this occurred by the fixation of two inversions that formed Strata 2 and 3 less than 5.9 and 4.7 million years ago, respectively, or 8.9 and 9.6 million years after the split with the blackspotted stickleback (Ross and Peichel 2008; Peichel et al. *in press*). In the blackspotted stickleback, the SDR expanded much more rapidly along Chr 19, with recombination suppressed between 10.5 and 14.1 million years ago, i.e., within 4 million years of the species split. As a result, the blackspotted Y is much more strongly degenerated than the threespine Y across much of the SDR. The boundaries of the SDR also differ, as the blackspotted PAR is much smaller (< 500 Kb) than the PAR in the threespine clade (2.5 Mb). Thus, Region 4 recombines in Japan Sea stickleback, but not blackspotted stickleback, further supporting that it evolved after the split with the threespine clade.

We identified three regions (Regions 2, 3, and 4) on the ancestral sex chromosome (Chr 19) of blackspotted sticklebacks that may represent distinct strata which ceased recombining at different times after the species split. They exhibit significantly different average values of *d*_S_, as expected of strata. However, extensive Y degeneration and resulting hemizygosity on Chr 19 in the blackspotted stickleback cause estimates of *d*_S_ to be unreliable. Also as expected of strata, these regions differ in the ratio of male vs. female read depth (Fig. 2). However, they also show clear differences in read depth in females, even though Y degeneration is expected only to reduce read depth in males (Fig. 2C). Region 3 contains the centromere (Sardell et al. 2018), which likely results in reduced read depth in both sexes. Region 4 also has much lower read depth in both the X and Y relative to the center of the chromosome. Reduced read depth towards the ends of Chr 19 is likewise present in Japan Sea sticklebacks (Dagilis 2019), and may be a sequencing artifact. Peichel et al. (*in press*) noted that the threespine stickleback PAR contains an abundance of transposable elements. If these elements evolve rapidly, sequences from closely related species may fail to map to this region of the threespine X reference. Unfortunately, previous studies of sex chromosomes in other species have relied solely on the read-depth ratio in males and females without presenting data from the sexes separately, so we cannot say how general this pattern is. Based on these findings, we suggest researchers not rely heavily on male-to-female read-depth ratios or *d*_S_ for defining strata when the differences between regions are small. Indeed, the distinction between Regions 1 and 2 cannot be detected using read depth ratio (Fig. 2) or *F*_ST_ (Fig. 3). The difference in *d*_S_ between them is statistically significant, but in the opposite direction of what is expected based on the known age of the regions relative to the species split. Instead, differences in gene tree topologies are the defining features of these strata (Fig. 6).

Population genetics statistics for the sex chromosomes differ between species as well. On Chr 19 of Japan Sea stickleback, the Y has lower molecular diversity than the X, while diversity is similar on the X and Y in the blackspotted stickleback. This difference is driven primarily by diversity on the X, which is much lower in blackspotted stickleback than Japan Sea stickleback (based on a similar pedigree-based study by Dagilis (2019)). This difference in diversity on the X may reflect differences in demography. The Japan Sea stickleback has undergone a major recent population expansion (Ravinet et al. 2018), while the blackspotted stickleback population appears to have recently contracted based on Tajima’s *D* for autosomal loci. Tajima’s *D* for the Y is strongly negative in both species, as expected since both Ys experienced a bottleneck and then a population expansion following the fusions that created their neo-Ys.

The ancestral Y has recently fused with an autosome in both the blackspotted and Japan Sea sticklebacks, but the identity of the autosome differs: Chr 12 is the neo-sex chromosome in the former, while Chr 9 is in the latter (Ross et al. 2009). Both these fusions occurred on the same end of Chr 19 distal to the PAR. Both are also very recent. We estimate that they occurred less than 1.4 million years ago in the blackspotted stickleback and less than 1.2 million years ago in the Japan Sea stickleback (Dagilis 2019) (Fig. 7). As a result, neither of these two neo-sex chromosome shows signals of extensive degeneration (see Kitano et al. 2009; Natri et al. 2013; Yoshida et al. 2014; Yoshida et al. 2017; Dagilis 2019). They do, however, differ greatly in the size of their SDRs. The nonrecombining region comprises 77% of the neo-sex chromosome in blackspotted sticklebacks but only 34% in the Japan Sea stickleback, even though the lengths of the fused chromosomes are nearly identical. Finally, in both species the neo-Y chromosomes have much lower molecular diversity than the neo-X and negative Tajima’s *D* (Dagilis 2019). These patterns may result from selective sweeps on the neo-Y associated with establishment of the fusion.

Why the blackspotted Y has much a much larger SDR and a much more highly degraded Y than the threespine clade is unknown. On one level, these differences likely reflect the ages of the non-recombining regions: it took more than twice as long after the species split for the inversion that formed Stratum 2 to fix in the threespine clade than it did for the SDR to expand across nearly all of Chr 19 in blackspotted stickleback. These differences may have arisen solely by chance.

Several factors are thought to also foster rapid rates of chromosome degeneration. These include higher mutation rates, shorter generation times, smaller population sizes, and increased reproductive skew among males (Graves 2006). Nothing is known about variation in mutation rates in sticklebacks, and all *Gasterosteus* species have similar generation times. The blackspotted stickleback has a much smaller global range than the threespine stickleback (Wootton 1976) and much lower molecular diversity on autosomes, which suggests it has a much smaller effective population size (based on comparisons with Hohenlohe et al. 2010). These differences may have made it easier for slightly deleterious inversions to drift to fixation on the blackspotted stickleback Y. Another possibility is that sexually antagonistic selection (SAS), which occurs when an allele is beneficial to one sex but harmful to the other, is stronger in blackspotted sticklebacks. Theory shows that SAS favors suppressed recombination between the X and Y and sex chromosome-autosome fusions (Charlesworth and Charlesworth 1980; Rice 1987; Charlesworth 2017). There is no *a priori* reason, however, to suspect that the degree of SAS should vary greatly between *Gasterosteus* species.

Another possibility is that differences in recombination landscapes between species could favor differential rates of SDR expansion. The strength of selection for inversions on sex chromosomes is directly proportional to the recombination rate between the sex determining gene and a locus subject to SAS (Rice 1987). The threespine clade of sticklebacks exhibits strongly sexual dimorphic recombination (*i.e*. heterochiasmy), with crossovers clustering near the telomeres of all chromosomes in males (Sardell et al. 2018). This pattern results in relatively low recombination between the X and Y throughout most of the chromosome. Low recombination, in turn, decreases the strength of selection for modifiers such as inversions that further reduce recombination between a site subject to SAS and the sex determining gene (Sardell and Kirkpatrick 2020). Strength of selection for SDR expansion might be stronger in blackspotted stickleback if the male recombination landscape is more uniform across chromosomes. Under this hypothesis, species differences in recombination landscapes would have evolved quickly since we estimate that the SDR began expanding in blackspotted stickleback very shortly after the species split. Although we do not currently have estimates of genome-wide recombination rates in blackspotted stickleback, this scenario is plausible as differences in fine-scale recombination landscapes rapidly evolved in very recently diverged populations of threespine stickleback (Shanfelter et al. 2019).

The conservation of Chr 19 as a sex chromosome for at least 14 million years, and likely 21.9 million years in *Gasterosteus* (Peichel et al. *in press*), contrasts sharply with the high rates of sex chromosome turnover in the rest of the stickleback family. Stickleback species in the genus *Pungitius*, for example, vary both in which chromosome is sex-linked and in which sex is heterogametic (Ross and Peichel 2008; Dixon et al. 2018; Natri et al. 2019). Why sex chromosome turnover rates vary so much within sticklebacks is a mystery. The “hot-potato” model of sex chromosome evolution posits that degeneration of the Y favors the invasion of a new Y chromosome in species without dosage compensation (Blaser et al. 2013). This hypothesis suggests that sex chromosome turnovers in *Gasterosteus* sticklebacks should be common since these species have degenerate Y chromosomes and incomplete dosage compensation (White et al. 2015). Perhaps some sort of constraint inhibits the origin of a new pair of sex chromosomes in *Gasterosteus* but not in other sticklebacks. Alternatively, SAS acting on Chr 19 may favor maintenance of the ancestral sex chromosome (van Doorn and Kirkpatrick 2007).

It is also unclear what evolutionary force drove the origin of the neo-sex chromosome of the blackspotted stickleback. SAS may have driven the fusion of Chr 12 in blackspotted stickleback, as data suggests may be the case in Japan Sea stickleback (Dagilis 2019). The targets of selection must differ, however, since the fusions involve different autosomes. Another possibility is the “fragile-Y” hypothesis (Blackmon and Demuth 2015). This theory posits that small PARs increase rates of aneuploidy in sperm because they provide little room for chiasma to form and ensure proper meiotic segregation. Fusions that expand the PAR may therefore be favored. Consistent with this idea, the PAR on Chr 19 is very small (0.4 Mb), comprising less than 2% of the total length of the chromosome. Moreover, its small size likely predates the fusion with Chr 12, a precondition of the fragile-Y hypothesis, as all regions of the Chr 19 SDR are much more degenerated than the Chr 12 SDR.

### Comparisons with other taxa

A handful of previous studies have also reported differences between closely related species in the extent of recombination suppression, sex chromosome differentiation, and Y (or W) degeneration. The best examples come from birds. Although all birds share an homologous W chromosome, the number of strata and size of the PAR varies widely between species (Zhou et al. 2014; Xu et al. 2019a; Xu et al. 2019b). Homologous strata also show extensive heterogeneity in degeneration across bird species (Zhou et al. 2014). Likewise, Hughes et al. (2005) observed much higher rates of degeneration on a part of the chimpanzee Y chromosome compared to its homologous region on the human Y. Both the avian and mammalian sex chromosomes are more than 140 million years old (Cortez et al. 2014). Our findings show that evolutionary dynamics of sex chromosomes can also differ dramatically between closely related species with much younger sex chromosomes (less than 22 million years old, based on Peichel et al. (*in press*)). Perhaps the most similar previous finding shows that heteromorphic and homomorphic sex chromosomes are homologous in the flowering plant genus *Spinacia* (Fujito et al. 2015). However, that study relied on cytogenetics, flow cytometry, and presence or absence of a small number of sex-linked markers to demonstrate differences in sex chromosome structure, and did not investigate the genomics of sex chromosome evolution. As such, they could only detect broad-scale variation in X-Y differentiation or Y degeneration across species.

Darolti et al. (2019) reported even more extreme variation in sex chromosome degeneration between closely related fishes. They showed that the Y chromosome of *Poecilia picta* is highly degenerate, even though the same pair of X and Y chromosomes are nearly undifferentiated in its congeners, *P. reticulata* and *P. wingei*. Darolti et al. (2019) did not explicitly establish that the Y chromosomes are homologous in all three species, however. Specifically, they did not rule out the possibility that a sex chromosome turnover occurred in the shared ancestor of *P. reticulata* and *P. wingei* (*i.e*., a new Y originated from an X chromosome after the ancestor of those two species diverged from *P. picta*). In this hypothesis, the sex chromosomes in the former two species are undifferentiated not because of slower rates of sex chromosome evolution, but because their Y chromosomes are much younger and/or continue to recombine. Darolti et al. (2019) found that surprisingly few *k*-mers are shared by all three species. They ascribed this result to extensive Y degeneration in *P. picta*. Yet we find shared sequences in the oldest stratum of the blackspotted and Japan Sea sticklebacks despite extensive Y degeneration in both species. Thus, their results are equally consistent with the turnover hypothesis, and this uncertainty precludes comparisons of rates of sex chromosome evolution. Unfortunately, the extreme degeneration of the *P. picta* Y chromosome imposes challenges for reconstructing the evolutionary history of these species’ sex chromosomes due to the difficulty in confidently assigning sequences to the Y.

For this study, we developed a novel method based on chromosome duplications that explicitly tests for the homology of sex chromosomes between species. This approach and a complementary method using gene trees are useful for comparing rates of sex chromosome differentiation and degeneration across closely related species. They also can be used to identify cryptic sex chromosome turnovers that involve the same linkage group. *Fugu* pufferfish and house flies (*Musca domestica*) are the only known cases of sex chromosome turnover events in which a new Y originated from an old X chromosome (Meisel et al. 2017; Ieda et al. 2018). Such turnovers may be much more common than currently believed, however, as it is much easier to identify turnovers that involve a transition between different pairs of chromosomes or between XY and ZW sex determination. We strongly recommend that all comparative studies of sex chromosome evolution explicitly test for cryptic turnover and Y chromosome homology. Not doing so can lead to underestimates of sex chromosome turnover and inappropriate comparisons between non-homologous Y chromosomes.

### Conclusions

Extensive variation in sex determination among stickleback fishes make them an ideal model system for studying sex chromosome evolution. Previous studies have documented several sex chromosome turnover events in this family (Ross et al. 2009; Dixon et al. 2018; Natri et al. 2019). The results of our study show that the situation in sticklebacks is even more complex, as the sex chromosomes of *Gasterosteus* sticklebacks differ substantially despite evolving from a common ancestral pair of XY chromosomes. We expect that future studies will show that similar variation within shared sex chromosome systems is common, and provide a novel framework for establishing sex chromosome homology.

## Materials and Methods

### Sampling and Sequencing

All procedures involving live fish were approved by the Veterinary Service of the Department of Agriculture and Nature of the Canton of Bern (VTHa# BE4/16 and BE17/17) and the St. Mary’s University Animal Care Committee (17-18A2). During June and July 2017, we collected threespine stickleback (*Gasterosteus aculeatus*) and blackspotted stickleback (*Gasterosteus wheatlandi*) from Canal Lake, Nova Scotia, Canada (44.498654, −63.902952), threespine stickleback from Humber Arm, Newfoundland, Canada (49.009842, −58.132643), and blackspotted stickleback (*Gasterosteus wheatlandi*) from York Harbour, Newfoundland, Canada (49.058555, −58.373138) under Department of Fisheries and Oceans permits for the Maritime Region (Licence 343930; FIN 700019217) and Newfoundland (Experimental Licence NL-4111-17). We made fifteen independent crosses by *in vitro* fertilization of the eggs of a single threespine stickleback female with sperm from a unique blackspotted stickleback male. Five crosses were made using five different blackspotted males and a single threespine female collected from Newfoundland, and ten crosses were made using ten different blackspotted males and three threespine females collected from Nova Scotia. Interspecific crosses were used because they allow us to better phase the paternal and maternal X chromosomes in daughters. These interspecies F1 hybrid embryos start to develop but then arrest and never hatch (Hendry et al. 2009; C. Peichel, pers. obs.), so crosses were raised until developmental arrest and then placed into 95% ethanol. DNA was extracted from fin clips of the cross parents and from individual embryos using phenol-chloroform extraction, followed by ethanol precipitation. The sex of the embryos was determined using microsatellite markers on chromosomes 12 (*Stn327*, *Pun2*) and 19 (*Stn284*, *Cyp19b*), that were previously shown to be sex-linked in blackspotted stickleback (Ross et al. 2009). For each of the fifteen crosses, we sequenced the mother, the father, one son, and one daughter. DNA from each of these individuals was used to construct Illumina TruSeq DNA nano libraries, which were sequenced for 300 cycles (2 x 150 bp paired-end reads) in an S2 flow cell on an Illumina NovaSeq 6000. Library construction, sequencing, barcode trimming, and initial quality control was performed by the University of Bern Next Generation Sequencing Platform.

### Sequence assembly & SNP calling

We used *bwa mem* v.7.16 (Li and Durbin 2010) to map all raw reads to the most recent masked assembly of the threespine stickleback reference genome (Glazer et al. 2015). We sorted and removed all reads with low mapping quality (< 20) using *SAMtools* v.1.6 (Li et al. 2009). SNP calling was performed using the *samtools mpileup* and *bcftools call* functions in *SAMtools* v.1.6 (Li et al. 2009). We used *VCFtools* v.1.15 (Danecek et al. 2011) to filter the VCF file, retaining only biallelic SNPs with minimum quality scores of 999. Genotypes where the genotype quality was less than 20 were treated as missing data. Finally, we removed all SNPs where more than 3 sons or 3 daughters were missing data.

We did not include the recently-published assembly of the threespine stickleback Y (Peichel et al. *in press*) in the reference genome used for mapping for several reasons. First, a main aim of this project was to identify homologous regions on the X and Y chromosomes that we could use to construct gene trees. This process would have been much more difficult if we had mapped reads to separate X and Y reference scaffolds. Second, much of the threespine stickleback Y has degenerated, especially in Stratum 1. We also expect that some genes that were lost on the threespine stickleback Y have been retained on the blackspotted stickleback Y. Thus, use of separate X and Y reference scaffolds would result in chimeric mapping of the blackspotted stickleback Y reads, with some aligning to the X reference and others aligning to the Y reference. This situation would greatly complicate phasing and our ability to directly compare the X and Y sequences.

### Phasing

We used a custom R script to phase the paternal and maternal gametes by transmission for parent-offspring trios, as described in Sardell et al. (2018) and Dagilis (2019). Briefly, for every heterozygous SNP in the offspring, we used parental genotypes to determine which allele was inherited from the father and which allele was inherited from the mother. Paternally inherited alleles in sons and daughters were transmitted in sperm containing an Y or X chromosome, respectively. This provided us with sequences of 15 blackspotted X chromosomes and 15 blackspotted Y chromosomes independently sampled from the wild. SNPs where the offspring and both parents were heterozygous cannot be unambiguously phased, so we conservatively treated them as missing data. Likewise, we removed any site where the offspring’s genotype included an allele that was not present in either parent. We then used a script provided by Dixon et al. (2018) to convert the genotypes into a haploid VCF file with a column for each gamete. We used custom phasing scripts rather than phase-by-transmission scripts from standard bioinformatic packages (e.g., GATK), because the latter are not specifically designed to account for transmission patterns in sex chromosomes and often produced clearly erroneous phasing results when applied to our data. Our approach is more conservative in assigning phased haplotypes to the X and Y.

### Identification of SDR

We used *VCFtools --geno-depth* v.1.15 (Danecek et al. 2011) to extract the read depth for each son or daughter at each SNP. We then used a custom R script to calculate the ratio of the mean read depths in sons to the mean read depths in daughters (i.e., read depth ratio) in 10 Kb windows. All custom scripts referred to in this section are publicly available, as outlined in the Data Accessibility section.

Before calculating population genetic statistics, we further filtered our genotypes using *VCFtools* v.1.15 to removes sites with excessive read depth, which likely represent duplications with multiple paralogs. We used 52 as the maximum mean read depth threshold, which represented 1.5 times the mean read depth on autosomes. We also removed sites where the mean read depth was below 26 (i.e., 0.75 times the mean autosomal read depth) to minimize genotyping errors at hemizygous sites. Finally, we used *VCFtools* v.1.15 to calculate weighted *F*_ST_ between the phased X and Y chromosomes, as well as genomic diversity (π) and Tajima’s *D* for the X and Y separately.

We calculated genomic divergence using a pipeline and scripts developed by Dixon et al. (2018). We first used FastaAlternateReferenceMaker from the Genome Analysis Toolkit (DePristo et al. 2011) to generate consensus sequences for the blackspotted X, blackspotted Y, and threespine X sequences from our pedigrees. We then used a custom script to generate individual sequences for each gene annotated in the .gff file for the most recent assembly of the threespine stickleback reference genome (Glazer et al. 2015). Finally, we used PAML v.4.9 (Yang 2007) to calculate *d*_S_, *d*_N_, and *d*_N_/*d*_S_ for each gene by comparing the blackspotted stickleback X and Y. We also calculated *d*_S_ by comparing the blackspotted stickleback X to the threespine stickleback X. We estimated the age of each region by taking the ratio of the mean synonymous site divergence between the blackspotted stickleback X and Y to the mean synonymous site divergence between the blackspotted and threespine stickleback Xs. We then multiplied the result by 14.3 million years, *i.e*., the estimated age of the most recent common ancestor between blackspotted and threespine stickleback (Varadharajan et al. 2019). We removed genes with fewer than 20 total SNPs when calculating *d*_N_*/d*_S_ on Chr 19 and fewer than 15 total SNPs on Chr 12, to eliminate division by zero errors. Lowering this threshold to 10 total SNPs did not affect the results significantly.

Gene trees within an SDR should be what we term “Y (or X) monophyletic” (Dixon et al. 2018; Dagilis 2019). This condition holds if all of the Xs or Ys form a monophyletic clade with respect to the other sex chromosome. We used a custom R script to convert the haploid VCF files for each chromosome into fasta files comprising the SNP genotypes for each individual. We then used RAxML v.8.2.12 (Stamatakis 2014), employing the GTRGAMMA model and a rapid bootstrap analysis (−*f a*) over 1000 bootstraps, to generate gene trees in 100 Kb windows across Chromosomes 19 and 12. We calculated the fraction of Ys (or Xs) falling within the largest monophyletic clade of Ys (or Xs) using a custom R script that employs the R packages *ape* (Popescu et al. 2012) and *phytools* (Revell 2012). Gene trees are considered to be Y monophyletic when all Y sequences fall within a single monophyletic clade.

### Identification of homologous autosomal-Y duplications

Our first approach for testing whether the sex chromosomes are homologous was to test whether chromosomal rearrangements involving the Y are shared between species. Bissegger et al. (2019) noted that, in threespine stickleback, many loci that map to autosomes in the Glazer et al. (2015) reference genome exhibit extreme differences in allele frequency between males and females. The observed allele frequency differences are biologically implausible as they require extreme mortality in the population (since autosomal allele frequencies will be approximately equal between males and females at conception). Bissegger et al. (2019) instead suggested that these regions represent loci that have duplicated from the autosome onto the Y chromosome. Alternatively, they may represent regions on the divergent Y chromosome that have duplicated onto an autosome and do not have a similar paralog on the X. The important result is that Y-linked reads erroneously map to an autosomal region of the reference genome because the reference was obtained from a female and the X does not contain an homologous sequence.

We remapped the blackspotted stickleback sequences from this study to the unmasked assembly of the Glazer et al. (2015) threespine stickleback reference genome (since repeat-rich regions may be more likely to duplicate). We again applied the same filtering criteria as those used for initial identification of the SDR. We removed any SNPs where we had high quality genotype data from fewer than 8 sons or fewer than 8 daughters. We did the same for a set of phased Japan Sea X and Y sequences generated by Dagilis (2019).

We tested for homology between the blackspotted and Japan Sea stickleback Y chromosomes by identifying autosomal regions that show extreme differences in allele frequencies between males and females in both species. We first used *VCFtools* v.1.15 to calculate weighted *F*_ST_ between the phased X and Y sequences separately for each species in 10 Kb non-overlapping windows across all autosomes. We then used custom R scripts to identify windows in which mean *F*_ST_ falls within the top 2 percent of windows in both species. Within each of these shared outlier windows, we calculated *F*_ST_ between the X and Y sequences at each SNP, and identified all SNPs with *F*_ST_ > 0.25 in both species. We further filtered the set of shared high-*F*_ST_ SNPs to include only those sites in which an allele is restricted to males in both species (as expected of mutations occurring on the Y paralog). Windows that contain multiple SNPs satisfying these criteria are interpreted as conclusive evidence that the Y chromosomes in blackspotted and Japan Sea sticklebacks are homologous. The alternative hypothesis of independent duplications would require not only that the same autosomal region was independently duplicated onto the Y multiple times, but also that it then independently accrued the same point mutations at multiple loci in less than 14 million years.

To confirm that the putative Y duplications are also present in threespine sticklebacks, we used *blastn* v.2.8.1 to search for similar sequences in the threespine stickleback Y reference (Peichel et al. *in press*) as well as the Glazer et al. (2015) threespine stickleback reference genome, which was sequenced from a female. We also identified significant overlap between autosomal regions with high *F*_ST_ from our study and the putative autosome-to-Y duplications identified by Bissegger et al. (2019) for threespine stickleback. We do not present the results in this manuscript, however, as the bioinformatic and analytic pipelines used in their study, including filtering criteria and measures of genetic differentiation, were quite different from ours, increasing the likelihood of type II errors.

### Tree-based method for identifying sex chromosome homology

Our second approach for testing whether the sex chromosomes are homologous in all three *Gasterosteus* species utilizes gene trees. We included phased blackspotted stickleback X and Y sequences from four crosses in this study. Each cross used a different threespine stickleback mother, and we included the phased maternal X sequences in our analysis. Four phased X and four phased Y sequences from Japan Sea stickleback fathers, and their eight corresponding phased maternal threespine X sequences, were obtained from an earlier study that employed the same experimental cross design (Dagilis 2019). SNP-calling and phasing for the combined data set was undertaken using the methodology described above. Any sites with mean read depth less than 17 or less than 67 (representing 0.5X and 2X mean read depth across all SNPs, respectively) were removed from the dataset. In addition, we removed any sites within the SDR that were heterozygous in the father but where phasing indicated that brothers and sisters inherited the same paternal allele. Such inheritance patterns cannot occur within an SDR and likely represent genotyping/phasing errors arising from hemizygosity on the Y. We rooted the trees using a computationally phased *Pungitius pungitius* genome from Dixon et al. (2018). Gene trees were constructed in RAxML v.8.2.12 (Stamatakis 2014) using the same parameters described above for testing XY consistency.

One potential problem with this approach is that genotyping errors in highly degenerate strata can lead to false topologies. For example, a SNP that is hemizygous in sons due to deletions on the Y will be assigned a homozygous genotype that incorrectly attributes the maternal threespine X allele to the blackspotted or Japan Sea Y. Therefore, we filtered the dataset to only include windows where the maximum likelihood tree either features four monophyletic clades representing the blackspotted Xs, blackspotted Ys, Japan Sea Xs, and Japan Sea Ys (as expected of old SDRs) or monophyletic clades for a species (as expected of PARs or new SDRs).

All windows within a non-recombining stratum should share the same topology since they all descend from the same physical chromosome on which recombination suppression (*e.g*., by an inversion) first arose. As described above, false topological inferences can arise from genotyping or phasing errors in regions of the chromosomes with large deletions on the Y. Therefore, we assume that the most common topology probably represents the true evolutionary history. Additionally, in our pedigrees, deletions on the Y are most likely to result in topologies in which one or both species’ Ys cluster with the threespine X sequences, since SNP-calling programs will wrongly impute the maternal threespine X allele as the blackspotted or Japan Sea Y genotype. Thus, topologies consistent with independent Y evolution (Fig. 5B) or Y turnover (Fig. 5C) are far more likely to arise via error than topologies consistent with a single Y origin (Fig. 5A).

## Data Accessibility

Sequencing data generated for this project are archived on the NCBI SRA database and will be released upon publication. Scripts used for data processing and analysis are available on GitHub (https://github.com/JasonSardell/BlackspottedStickleback). Scripts and intermediate data files are also archived on Dryad and will be released upon publication.

## Acknowledgments

This work was supported by the National Institutes of Health (grant R01-GM116853 to M.K. and C.L.P.). We thank Andrius Dagilis for assistance on genomic analysis and comments on the manuscript, and Groves Dixon for providing data and computer scripts.

---

**Supp. Fig. S1:**
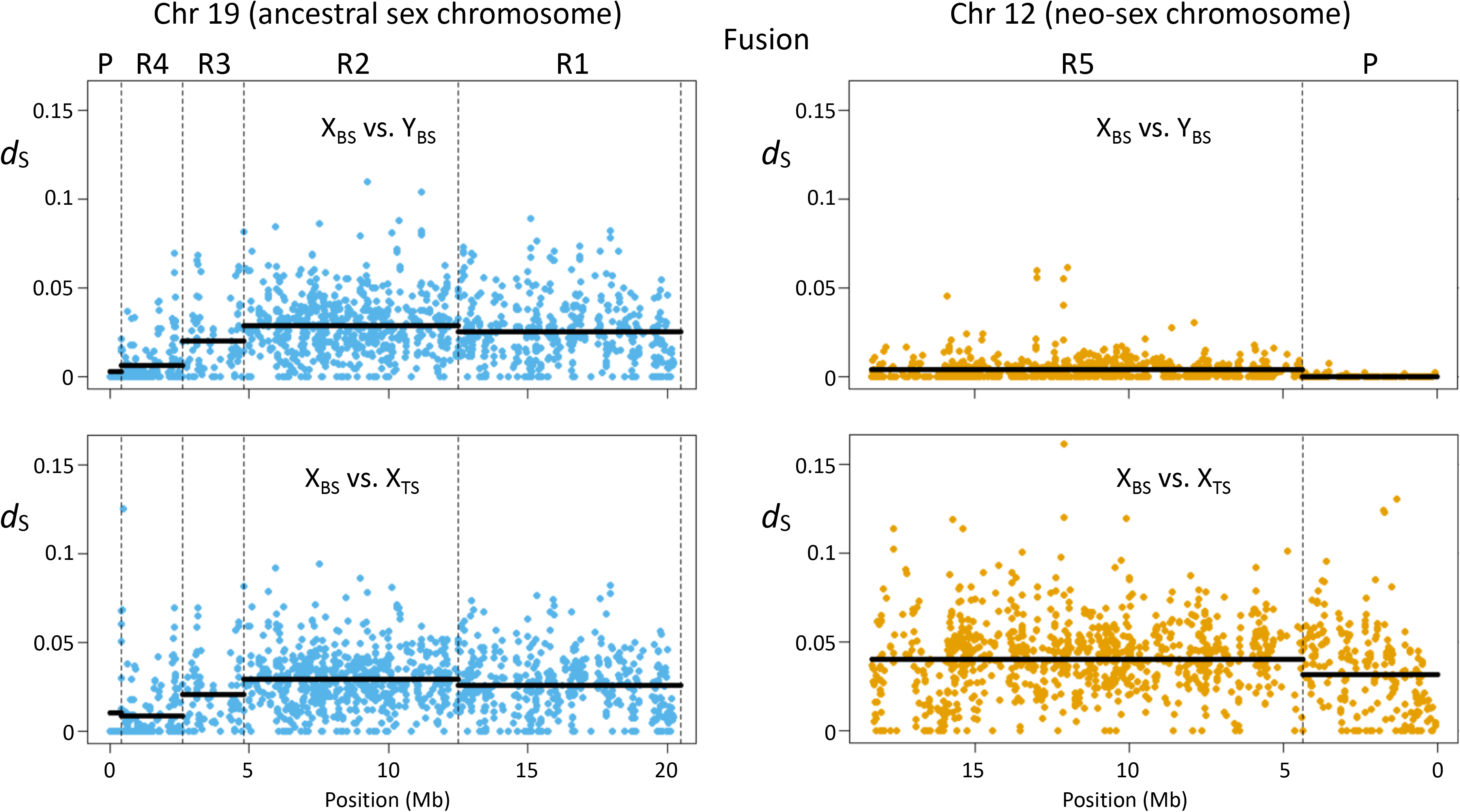
Divergence at synonymous sites (*d*_S_) for each gene on Chr 19 (left panels) and Chr 12 (right panels). Top panels: *d*_S_ between blackspotted stickleback (BS) X and Y chromosomes. Bottom panels: *d*_S_ between blackspotted stickleback (BS) and threespine stickleback (TS) X chromosomes. Dashed vertical lines represent boundaries between the PARs (labeled P) and the four regions in the SDR on Chr 19 (labeled R1 to R4) and on Chr 12 (labeled R5) that were identified using methods described in text. Solid horizontal lines show the means for the regions.

**Supp. Fig. S2:**
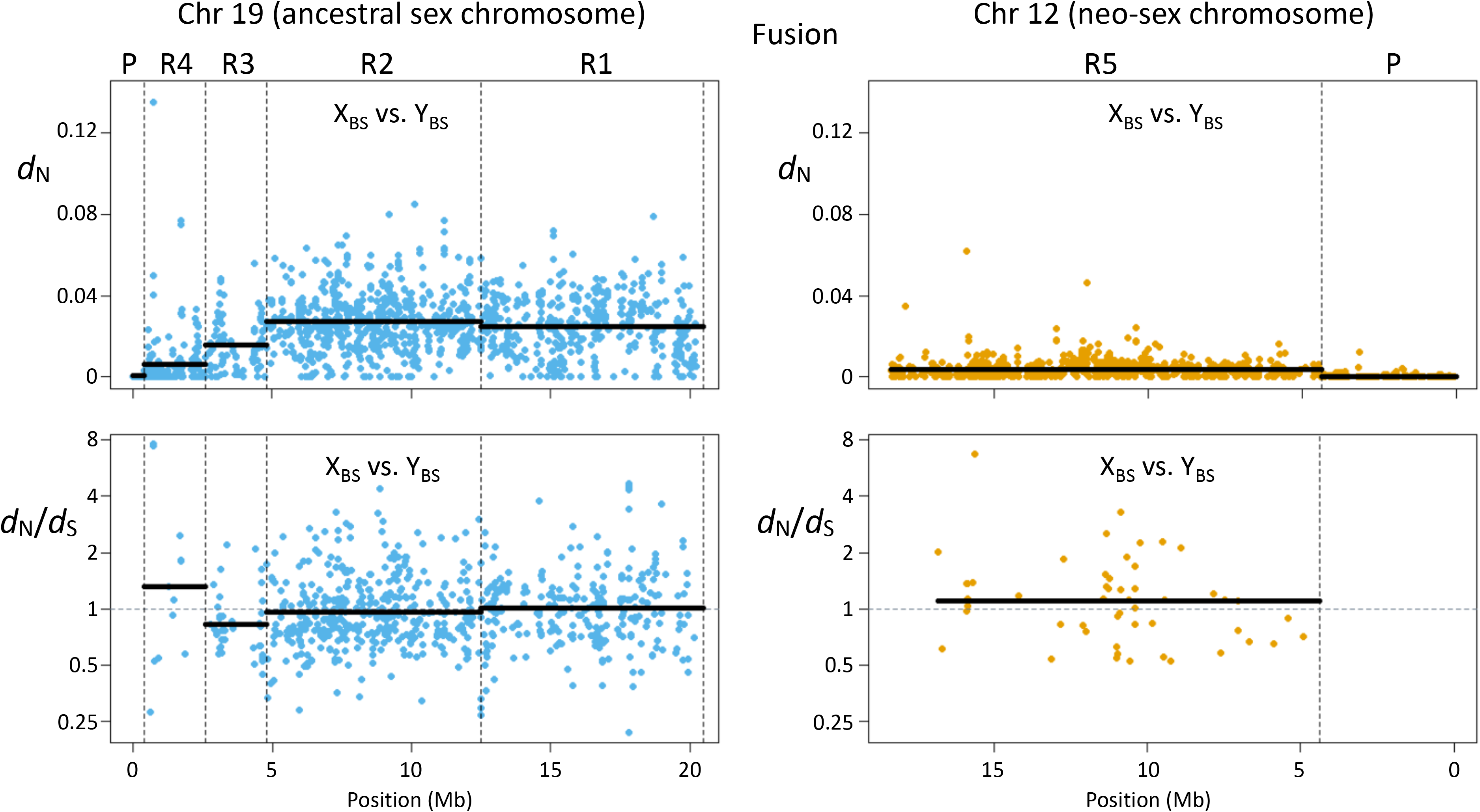
Divergence statistics for individual genes on Chr 19 (left panels) and Chr 12 (right panels). Top panels: divergence at nonsynonymous sites (*d*_N_) between blackspotted stickleback (BS) X and Y chromosomes. Bottom panels: *d*_N_/*d*_S_ between blackspotted stickleback X and Y chromosomes. Only genes containing at least 20 SNPs (Chr 19) or 15 SNPs (Chr 12) are included in *d*_N_*/d*_S_ plot to eliminate division by zero errors. Dashed vertical lines represent boundaries between the PARs (labeled P) and the regions in the SDR on Chr 19 (labeled R1 to R4) and on Chr 12 (labeled R5) that were identified using methods described in text. Solid horizontal lines show the means for the regions. The Y axes are plotted on a log scale.

**Supp. Fig. S3:**
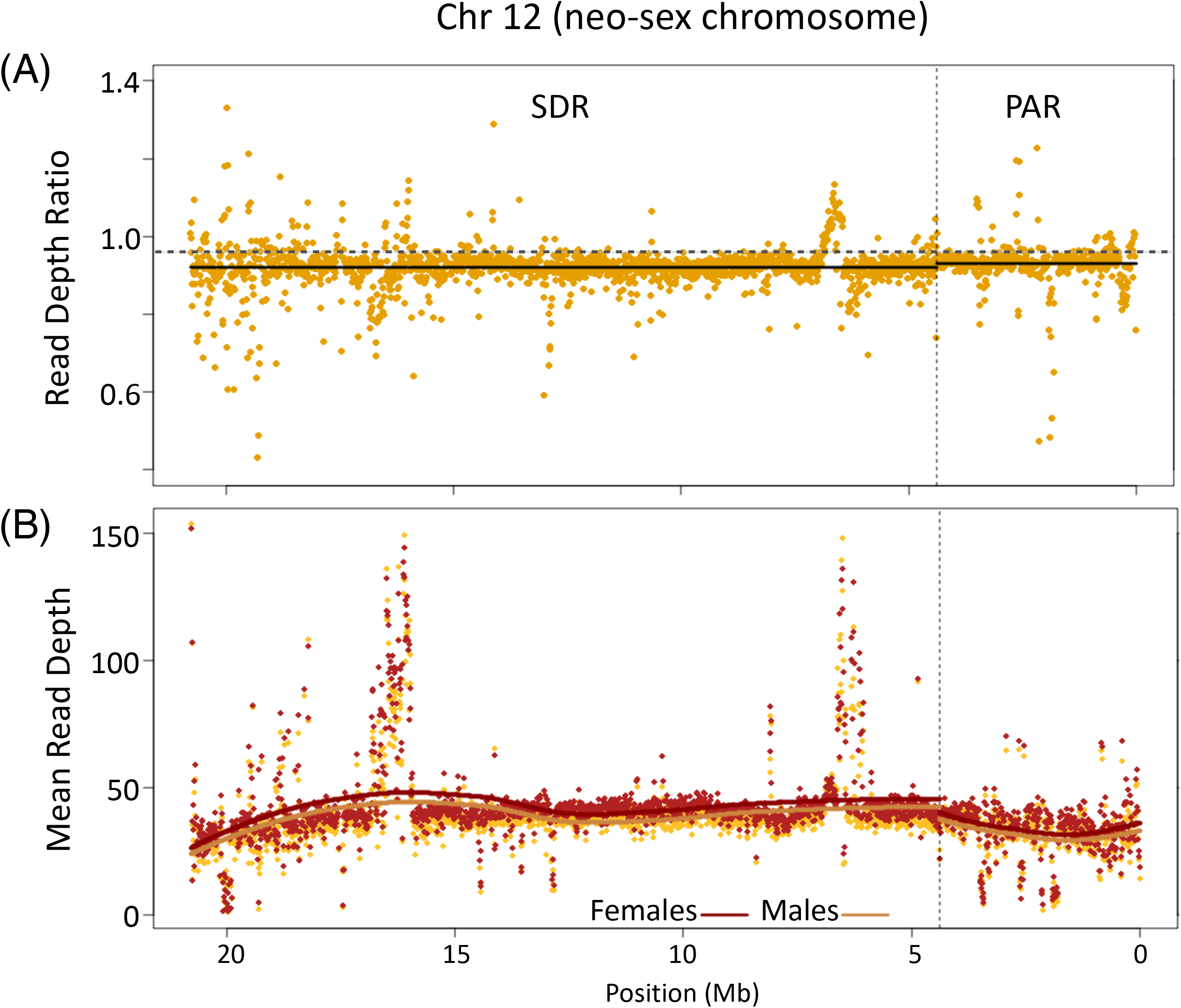
Read depth statistics for Chr 12 of blackspotted stickleback based on 15 sons and 15 daughters. (A) Plot of mean male/female read depth ratio. Dashed horizontal gray line represents the autosomal mean. (B) Plot of mean read depth in sons (orange) and daughters (red). Dots show averages in 10 Kb windows. Dashed vertical lines show the boundary between the SDR and the PAR. Solid curves are Loess best-fit.

**Supp. Fig. S4:**
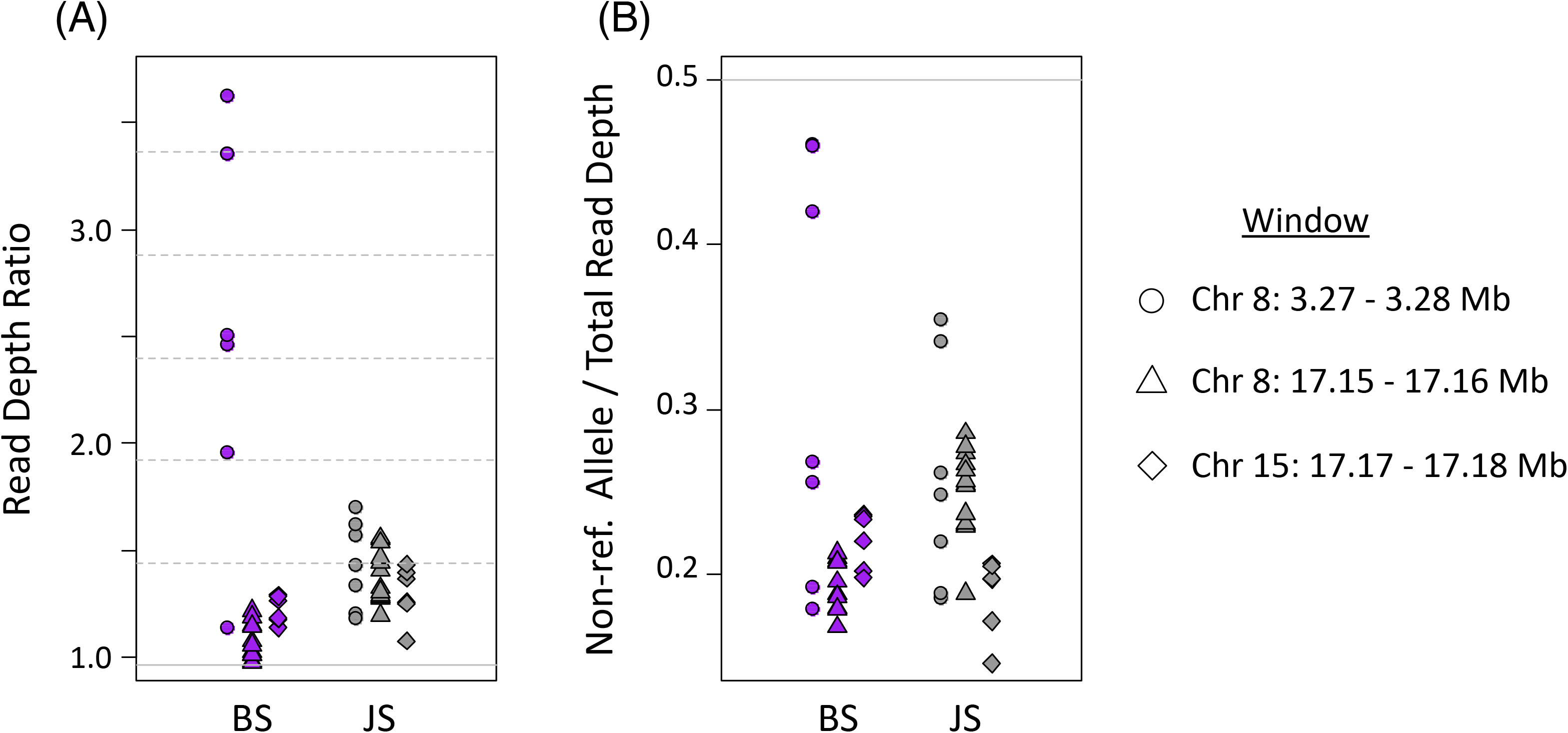
Read depth statistics provide evidence for autosome-to-Y duplications in the three regions of the blackspotted SDR shown in Fig. 4. Each point represents a SNP with *F*_ST_ > 0.25 and a male-specific allele in both blackspotted and Japan Sea sticklebacks. Points denote SNPs from the 10 Kb windows with the shapes shown in the key. (A) Male/female read depth ratio in blackspotted (BS) sticklebacks (left, purple) and Japan Sea (JS) sticklebacks (right, gray). Solid horizontal gray line indicates the autosomal mean. Horizontal dashed gray lines indicate intervals of 0.5 times the autosomal mean, i.e., the expected values for one or more autosome-to-Y duplications. (B) Fraction of reads possessing male-specific allele in blackspotted (BS) sticklebacks (left, purple) and Japan Sea (JS) sticklebacks (right, gray). Solid horizontal gray line indicates the value expected for non-duplicated regions.

**Supp. Fig. S5:**
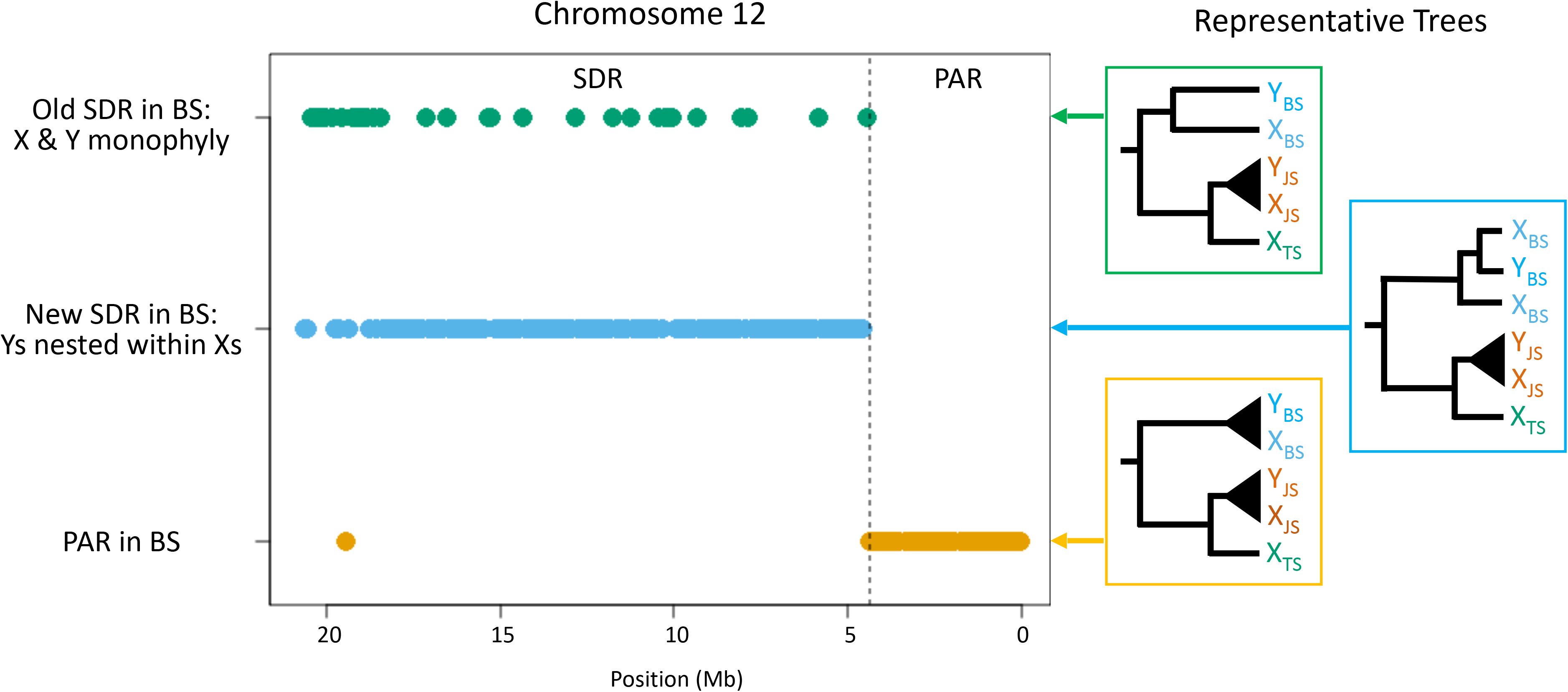
Gene tree topologies along Chr 12 demonstrate the recent origin of the SDR. Each dot represents the maximum-likelihood topology for a 100 Kb window. Representative trees are shown at right (BS = blackspotted, JS = Japan Sea, TS = threespine). In most windows from the SDR, a monophyletic clade of blackspotted stickleback neo-Ys is imbedded within the blackspotted neo-Xs.

